# Immersion in nature attenuates the development of mechanical secondary hyperalgesia: a role for insulo-thalamic effective connectivity

**DOI:** 10.1101/2024.10.11.617804

**Authors:** Sonia Medina, Sam Hughes

## Abstract

Nature-based social prescribing has been shown to improve physical and mental health and is increasingly used to manage chronic pain using immersive virtual reality (VR). However, the mechanisms of nature-based analgesia during immersive VR experiences remain unclear. In this study, we used experimentally induced sensitisation within central nociceptive pathways using high frequency stimulation (HFS) over the right forearm in 30 healthy participants and tracked the development of secondary hyperalgesia across three conditions: immersive VR nature, non-immersive 2D nature video, and no intervention. Immersive nature VR significantly reduced the development and spread of hyperalgesia, with sustained analgesic effects correlating with perceived presence. Bayesian modelling of neuroimaging endpoints collected separately revealed nature VR induced analgesic effects correlated with insulo-thalamic effective connectivity. We propose that the analgesic effects of nature are likely mediated via top-down endogenous analgesic systems which could be working to reduce the development and spread of heterotopic plasticity in the spinal cord.

## Introduction

There is a growing interest in the relationship between nature and pain relief. Therapeutic effects of nature exposure or nature-based social prescribing have been demonstrated in several chronic pain conditions [1–4], which could help to reduce the overall global burden of pain [5]. Nevertheless, it is still unclear how nature exerts analgesic effects. Recent theoretical frameworks for nature-based analgesia point towards potential activation of endogenous analgesic systems through top-down autonomic and/or cognitive processes [6]. However, the exact mechanisms are still poorly understood.

The rapid development of immersive virtual reality (VR)-based technologies has provided new ways to understand biopsychosocial pain mechanisms [7]. Currently, it is possible to investigate neural and psychophysical mechanisms of highly realistic, real world, nature-based immersive 360-degree experiences in a controlled lab-based setting [8]. Certain features of nature are thought to contribute towards therapeutic effects; the sense of being away, a good person-environment fit, sufficient visual engagement or features that can capture attention effortlessly, and should be considered when designing nature-based VR-based investigations [6]. It is therefore key to use natural environments which score highly on these qualities, a concept related to perceived restorativeness [9]. It is also important to use adequate non-immersive nature control conditions with a different level of perceived presence, which is associated with having different levels of both immersion and realism, to the immersive nature VR condition [10, 11].

By employing nature-based VR experiences alongside the high frequency stimulation (HFS) model of central sensitisation in healthy participants [12], it is possible to explore the effects of nature-based experiences on the development of mechanical secondary hyperalgesia, which mimics clinically relevant phenotypes seen during neuropathic pain [13]. Moreover, by acquiring resting-state functional magnetic resonance imaging (fMRI) from the same participants, we could potentially study how supraspinal models of endogenous pain modulation within Bayesian predictive frameworks [14–17] relate to the variability in analgesic effects between immersive and non-immersive nature exposure. Such an approach may help us explore whether, and by which mechanisms, nature-based VR may provide more effective pain relief than non-immersive nature interventions.

In this study, we investigated the effects of immersive and non-immersive virtual nature exposure on the development of mechanical secondary hyperalgesia using the HFS model in healthy participants, compared to a no-intervention control condition. Furthermore, we explored the underlying neurobiological mechanisms of these responses through dynamic causal modelling analysis on fMRI data at rest. Our preregistered hypotheses were:

1. Exposure to VR-nature during central nociceptive pathways sensitisation by HFS would attenuate the development of secondary hyperalgesia.
2. There would be relationships between cortical and sub-cortical resting state effective connectivity and responses to VR-nature exposure.
3. Modulation of pain responses by VR-nature will be correlated with the degree of perceived restorativeness and presence by each participant.
4. Restorativeness and presence will be higher for the VR-nature condition compared to non-immersive nature condition.

## Methods

### Participants

Thirty healthy, pain-free volunteers participated in this study (18 female; mean age = 29 years, SD = 10). Participants were screened to ensure no history of neurological or psychiatric disorders, acute migraines with loss of consciousness, dizziness, or motion sickness. Additional exclusion criteria included substance or alcohol abuse, inability to lie flat still, use of medications affecting temperature sensitivity or endogenous analgesia, cardiovascular drugs, inability to understand instructions, and MRI contraindications (e.g., metal implants, pacemakers, pregnancy). Participants were instructed to abstain from alcohol and recreational drugs 24 hours prior to each visit, avoid painkillers or paracetamol for 12 hours before sessions, and limit to one caffeinated drink in the morning. Compliance with these guidelines was confirmed at the start of each visit, and any deviations led to immediate termination and rescheduling. Informed consent was obtained from all participants at the beginning of their first visit. The study was approved by the Health Research Authority and Health and Care Research Wales ethics committee (Ethics reference: 22/HRA/4672). Participants could withdraw from the study at any time.

### Experimental design

Participants attended four study visits. For visits 1-3, a randomised crossover design was followed with three conditions: ‘VR’, ‘sham-VR’ and ‘no intervention’. The fourth visit consisted of an fMRI session (Figure 1). At the start of each session, compliance with lifestyle guidelines was assessed. Participants also completed the state part of the State Trait Anxiety Inventory[18] and were instructed to answer the question ‘are you in any pain today?’ on a numerical rate scale (NRS) ranging from 0 (‘no pain at all’) to 100 (‘wors pain imaginable’). Visits 1-3 were scheduled at around the same time of the day for each participant and around one week apart. The MRI visits was scheduled at least 2 days apart from the other visits.

**Figure 1.**
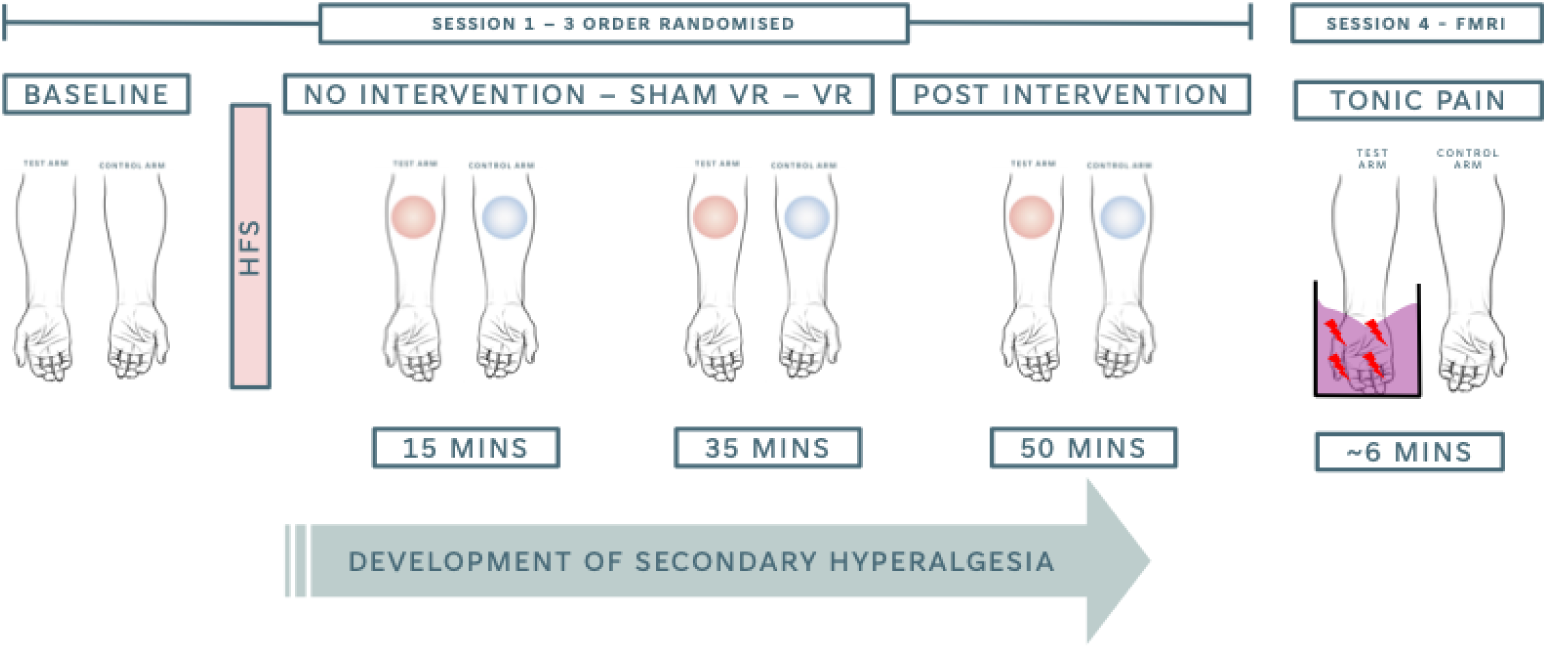
Experimental design. HFS = high frequency stimulation; fMRI = functional magnetic resonance imaging. Red circles represent mapping area following HFS. Blue circles represent mapping area for the control arm (i.e., no HFS). For sessions 1-3, arm testing order was counterbalanced across participants but remained consistent within participants. Session 4 also included a T1 anatomical scan (not depicted here) and one more resting state scan before and after the tonic pain condition (not included in the current manuscript). HFS = high frequency stimulation; FMRI = functional magnetic resonance imaging.

### Psychometric psychological screening

Participants completed the following psychological screening questionnaires: the Revised Cognitive Coping Strategies Inventory (CCSI-R)[19]; the Eysenck Personality Questionnaire-Revised Short Scale (EPQ-R)[20]; the trait part of the State Trait Anxiety Inventory (STAI)[18].

### High-frequency stimulation model (visits 1-3)

Electrical stimuli were administered to the centre of the right volar forearm using an epicutaneous pin electrode consisting of a circular array of 15 cathodal pins (individual diameter: 0.2 mm; length: 1 mm; overall diameter: 10 mm; area: 79 mm²), surrounded by a stainless-steel anode (inner diameter: 20 mm; outer diameter: 40 mm). A constant current stimulator (pulse width: 2 ms; DS7; Digitimer Ltd, Welwyn Garden City, UK) delivered each stimulus. Participants’ forearms were cleaned with isopropyl alcohol prior to electrode placement. The electrode’s location was defined by delineating a circular region matching its diameter, marked with four landmarks 1 cm apart along eight radial axes. An identical grid was drawn on the left volar forearm for consistency in control assessments. Each visit began with determining the individual electrical detection threshold (EDT) using the method of limits [21]. Pulses were applied starting at 0.05 mA in 0.05 mA increments until participants reported a clear sensation, then decreased by 0.01 mA until the sensation was lost. The intensity was increased in 0.01 mA increments until the sensation returned, repeating this process three times. The average of the six recorded intensities represented each participant’s EDT. For high-frequency stimulation (HFS), five trains of 100 Hz stimuli were controlled via a pulse generator (D4030; Digitimer Ltd) lasting one second, with ten-second intervals. These trains were delivered at 20 times the individual EDT. Participants rated the intensity of HFS following each stimulus on a numerical rating scale (NRS).

### Mechanical pain sensitivity assessments (visits 1-3)

Participants’ mechanical pain sensitivity (MPS) across both forearms was assessed using a shortened version of the DFNS MPS protocol. Three different pinprick stimuli (128 mN, 256 mN, and 512 mN) were applied five times in pseudo-randomized order within a 1 cm area around the electrode. Stimuli locations were alternated across four quadrants to prevent windup effects. Following each stimulus, participants rated their perceived pain on a numerical rating scale (NRS). MPS assessments occurred at baseline, and at 15, 35, and 50 minutes post-HFS. After each MPS assessment, the previously delineated landmarks were stimulated with a 256 mN pinprick, and participants rated their pain on an NRS to map secondary hyperalgesia (Figure 2). The order of arm stimulation was counterbalanced across participants but consistent within individuals. Participants kept their eyes closed during all baseline and post-intervention assessments, as well as the ‘no intervention’ session. During VR and sham-VR interventions, participants were instructed to keep their eyes open, looking straight ahead.

**Figure 2.**
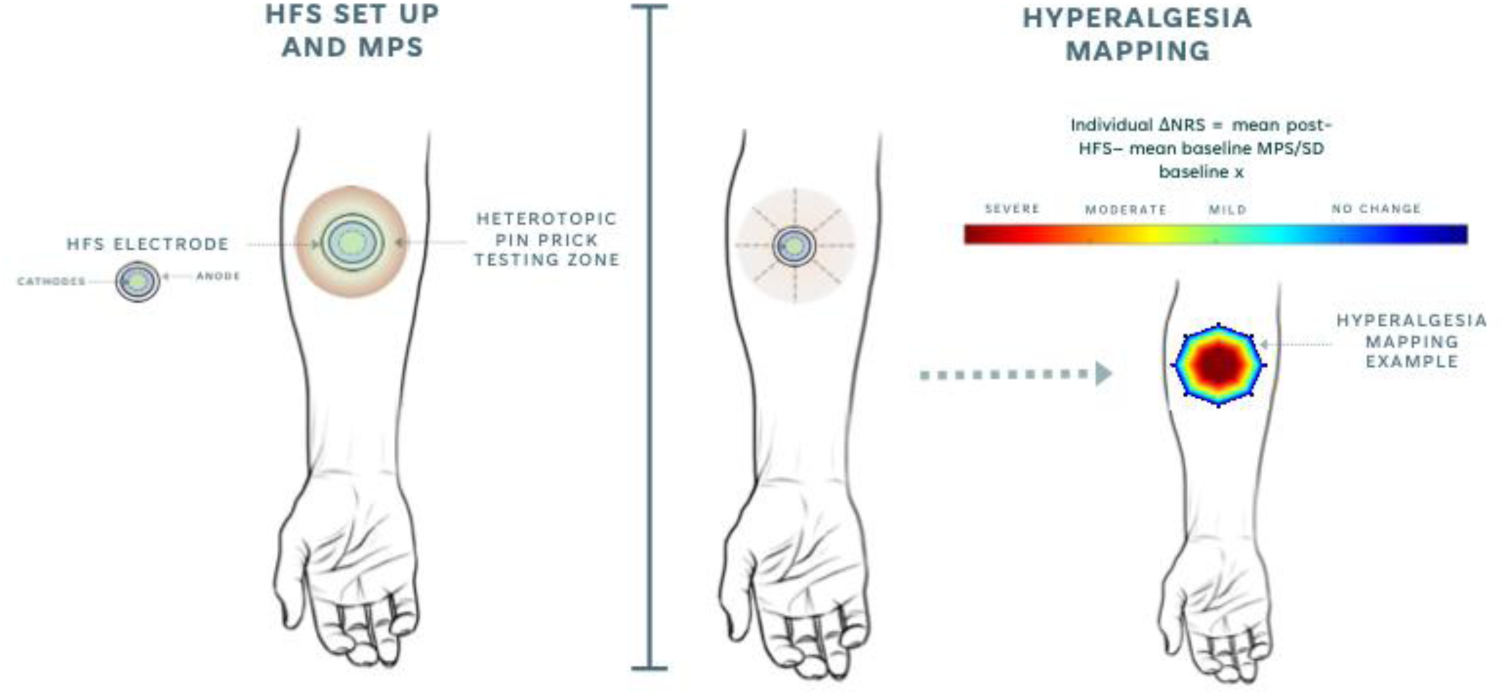
HFS set up (left) and hyperalgesia mapping procedure and calculation (right). prior to the placement of the electrode and the beginning of psychophysiological testing, participants’ forearm surfaces were cleaned, and an eight-radial axes grid was drawn on both forearms, approximately in the same equivalent position. HFS electrode was removed from participants’ skin immediately after HFS was performed. MPS = mechanical pain sensitivity; NRS = numerical rating scale; SD = standard deviation.

### Interventions (visits 1-3)

#### No intervention

Following HFS and in between MPS assessments, participants were asked to complete all the psychometric questionnaires in the same order. Participants who finished before the 50-minutes post HFS MPS assessment were instructed to stay seated and in silence for the remaining time.

#### VR intervention

Following HFS in the VR visit, participants completed 45 minutes of immersion in a nature scene with visual and auditory inputs through an HTC Vive Pro 2 headset. The scene consisted of a 4k footage of the Trail of the Ten Falls in Oregon, USA, available in Youtube (https://youtu.be/HvsBCN34dKY?si=7baBhh6-aCbWOVmk). During the intervention, participants were instructed to remain seated and were allowed to freely explore the 360-degree scene on a rotating chair.

#### Sham-VR intervention

Following HFS in the sham-VR visit, participants watched the same footage as the one presented in the VR session on a 2D screen while the room was kept otherwise quiet and with deemed lights. Participants were given the opportunity to explore the full 360-degree scene with the keyboard.

### Psychometric assessment of intervention

Following VR and sham-VR interventions, and before the 50-minute post HFS assessment, participants completed the Perceived Restorativeness Scale [9] and the Presence Questionnaire [22], in order to characterise differences in individual experiences regarding the nature scene and the VR/sham-VR setup.

### Functional MRI (visit 4)

Data were collected using a 3T Siemens Prisma MRI scanner equipped with a 64-channel. Participants underwent T1-weighted structural scanning (repetition time (TR) = 2100 ms; echo time (TE) = 2.26 ms voxel size = 1 x 1 x 1 mm) and a ∼6 minutes multi-echo, multi-band echo planar imaging (EPI) scan (TR = 1500 ms; TE1= 11 ms, TE2 = 27.25 ms, TE3 = 43.5 ms , TE4 = 59.75 ms; 51 slices with a thickness of 2.60 mm; voxel size = 2.6 x 2.6 x 2.6 mm; field of view 220 mm2, flip angle 77°; 229 volumes; total scanning time = ∼6 minutes). Slices were acquired in interleaved order. In addition, a field map scan was acquired at the end of the functional scan (TR = 590 ms; TE1= 4.92 ms, TE2 = 7.38 ms; 51 slices with a thickness of 2.60 mm; voxel size = 2.6 x 2.6 x 2.6 mm; field of view 220 mm2, flip angle 46°; interleaved acquisition). fMRI scanning reported here was acquired while participants immersed their right hand in a noxious cold gel, with the purpose of eliciting a tonic painful sensation. A full report of materials, procedures and preprocessing from the full imaging session can be found in the Supplementary Information (a).

### Data analysis

#### Individual experiences during VR and sham-VR interventions

To explore differences in restorativeness and presence measures between VR and sham-VR, a two-way repeated measures analysis of variance (ANOVA) was conducted with two within-subjects factors: Questionnaire (PRS and PQ) and Intervention (VR and sham-VR). Differences within each questionnaire across sessions were examined using separate paired-samples t-tests.

#### Secondary hyperalgesia effects

MPS measures were defined as the geometric mean of the 15 NRS scores from pinprick stimulation. To assess changes in MPS after HFS on each arm, we computed the percentage change from baseline MPS to each post-HFS assessment. MPS changes from baseline within each session were z-transformed as follows: standard ΔMPS = (MPSindividual − MPSbaseline) / SDbaseline. The effect of interventions on percentage change in MPS and standard ΔMPS within each session was assessed using one-way repeated measures ANOVA with Time as a factor with four levels (baseline, 15, 35, and 50 minutes post-HFS). Identical analyses were performed for each pinprick intensity (128 mN, 256 mN, 512 mN). Interaction effects between conditions and timepoints were explored via a two-way repeated measures ANOVA with two within-subjects factors: Time (baseline, 15, 35, 50 minutes post-HFS) and Condition (no intervention, VR, and sham-VR). Results were considered significant at p < 0.05. To examine the extent of secondary hyperalgesia, the percentage change and standard change from baseline NRS scores from 256 mN pinprick stimulation across eight axes were computed for each arm, session, and timepoint separately.

#### Relationships between individual experience during interventions and secondary hyperalgesia post-intervention

To explore whether individual differences in perceived restorativeness and presence across interventions related to lingering effects of secondary hyperalgesia post-intervention, we calculated Pearson’s correlation between delta scores across sessions within each questionnaire (sham-VR – VR) and MPS results 50 minutes post-HFS (sham-VR – VR). This represents a deviation from the preregistered analysis plan, where correlations between questionnaires and MPS results were initially planned for the 35-minute post-HFS timepoint. This decision was based on the fact that at 35 minutes post-HFS, participants were still receiving the intervention, so a true relationship between PRS and PQ and the lingering effects of the interventions could not be established.

All behavioural statistical analyses were performed on IBM SPSS Statistics Software Package v29.0, with initial computations (i.e., percentage of change, ΔMPS) performed in Microsoft Excel.

#### Effective connectivity analysis

We hypothesised that the analgesic effects of nature-based VR immersion might be mediated by enhanced parasympathetic tone through improved autonomic regulation. To investigate this, we set out to explore whether participants who showed greater engagement of brain circuitry associated with descending pain control during tonic noxious stimulation would experience more pronounced analgesic effects from nature-based VR. This hypothesis aligns with the idea that the effectiveness of our nature-based VR intervention could be linked to the basal ability of certain brain networks (particularly those involved in autonomic regulation and top-down modulation of pain) to respond to and modulate pain. We carried out spectral dynamic causal modelling (spDCM) to determine whether the patterns of effective connectivity (i.e., the excitatory or inhibitory influence of one brain region on another) within an a priori defined network could account for variability in net effect of the VR intervention in MPS at 50-minutes post intervention. The network was formed by seven a priori regions of interest (ROI): the dorsal anterior cingulate cortex (dACC) bilaterally, the anterior insula (AI) bilaterally, the thalamus bilaterally and the periaqueductal grey (PAG). Full details on ROI description can be found in the Supplementary Information (b).

We carried out all spDCM within the DCM toolbox implemented in SPM12 [23]. First, a fully connected model considering all possible connections across ROIs was specified for each participant. These models were then inverted using the variational Laplace scheme, which estimates model parameters by optimising the balance between data fit and model complexity, ensuring a trade-off between explanatory power and parsimony. For the inversion, we utilised the default probability densities provided by SPM12. At group level, individual differences were modelled using parametric empirical bayes (PEB) [24]. The design matrix included the group mean, and the net effect of the VR intervention in MPS measures 50-minutes post HFS, computed as follows: net VR effect = *MPS_50min_ following VR* − *MPS_50min_ following no intervention*. We applied default probability densities provided by SPM12. Given the absence of a specific hypothesis regarding the most plausible models, we conducted an automatic search over nested PEB models using Bayesian model reduction (BMR) [24]. The final model parameters were derived by averaging the values from the 256 models generated in the last iteration of the algorithm, with each model weighted by its evidence through Bayesian model averaging (BMA) [25]. Only parameters with a posterior probability greater than 95% of being present versus absent are reported.

#### Correlation with behavioural measures

To further interpret the relationship between parameters resulting from BMA and net VR effect, we carried out canonical variate analysis (CVA) and we computed Pearson’s correlation between the maximum contributing parameter to the canonical variate and the net VR effect.

## Results

### Participants

One participant did not report any of the tests performed and was therefore excluded from final data analysis (final N=29). One participant did not complete the fMRI session due to anxiety inside of the MRI scanner (final spDCM N=28). One participant reported mild headache five minutes before the end of the VR scene, and intervention was therefore terminated early. No other side effects were reported. All psychometric screening results can be found on Table 1.

**Table 1.** Psychometric assessment results. CCSI-R = cognitive coping strategy inventory; EPQ = Eysenck personality questionnaire; STAI = state and trait anxiety inventory; PRS = perceived restorativeness scale; PQ = presence questionnaire; BA = being away; SD = standard deviation.

| Measure | Screening | VR | Sham-VR | t(28) | sig |
| --- | --- | --- | --- | --- | --- |
|  | Mean (SD) | Mean (SD) | Mean (SD) |  |  |
| CCSI-R Distraction | 2.38 (0.73) | - | - | - | - |
| CCSI-R Catastrophising | 2.27 (0.79) | - | - | - | - |
| CCSI-R Minimisation | 3.12 (0.57) | - | - | - | - |
| EPQ Extraversion | 13.52 (5.29) | - | - | - | - |
| EPQ Neuroticism | 14.00 (6.77) | - | - | - | - |
| EPQ Psychotism | 5.45 (3.18) | - | - | - | - |
| EPQ Lie | 7.34 (3.53) | - | - | - | - |
| STAI Screening state | 32.24 (10.87) | - | - | - | - |
| STAI_Screening trait | 41.86 (11.96) | - | - | - | - |
| STAI State | - | 29.69 (8.63) | 24.55 (13.63) | 1.887 | 0.070 |
| PRS BA | - | 18.93 (7.76) | 17.83 (7.20) | 1.00 | 0.324 |
| PRS Fascination | - | 32.86 (10.98) | 30.41 (12.75) | 1.58 | 0.124 |
| PRS Coherence | - | 19.41 (4.20) | 21.03 (2.47) | -2.52 | 0.018* |
| PRS Compatibility | - | 31.83 (10.55) | 30.76 (12.03) | 0.59 | 0.559 |
| PRS Total | - | 103.03 (26.21) | 100.03 (27.14) | 0.80 | 0.428 |
| PQ Realism | - | 30.10 (8.38) | 23.31 (8.30) | 4.59 | 0.001** |
| PQ Possibility to Act | - | 13.90 (5.77) | 12.93 (5.95) | 0.86 | 0.396 |
| PQ Quality of Interface | - | 14.24 (3.05) | 14.21 (3.04) | 0.04 | 0.963 |
| PQ Possibility to examine | - | 10.45 (3.59) | 8.79 (3.19) | 2.27 | 0.031* |
| PQ Self evaluation | - | 10.34 (2.66) | 9.14 (3.37) | 1.80 | 0.083 |
| PQ Sounds | - | 15.66 (3.77) | 13.76 (3.87) | 2.73 | 0.011* |
| PQ Haptic | - | 2.52 (1.82) | 2.34 (1.08) | 0.42 | 0.677 |
| PQ Total | - | 97.21 (17.79) | 84.48 (18.39) | 3.83 | <0.001** |

### Perceived restorativeness and presence during VR and sham-VR

Two-way repeated measures ANOVA revealed a significant effect of Questionnaire (F_(1,28)_=9.877, *p*=0.004) and Intervention (F_(1,28)_=7.274, *p*=0.012), as well as significant interaction effect (F_(1,28)_=5.966, *p*=0.021). Paired t-tests within each questionnaire indicated that while there was not a significant difference in total perceived restorativeness, the presence scores were significantly higher during the VR intervention than during the sham-VR intervention (Table 1).

### Effect of HFS in mechanical pain measures

*Percentage of change*: Within the test arm, two-way repeated measures ANOVA revealed a main effect of Timepoint (F_(2.56)_=6.098, *p*=0.012). There were no significant main effects of Condition or Timepoint x Condition interactions (Table 2). One-way ANOVAS revealed a main effect of Timepoint during the no intervention condition (F_(3,84)_=8.495, *p*<0.001), with post-hoc pairwise comparison reflecting significant increase of MPS measures from baseline at all timepoints of over 130%. During the VR and sham-VR conditions there were no significant changes in MPS observed. On the control arm, there were no main effects of Condition, Timepoint or Condition x Timepoint interactions. One-way ANOVA’s indicating a significant effect of Timepoint only for the VR condition (F_(3,84)_=3.000, *p*=0.047), derived from a significant 25.79% decrease in MPS measures at 35-minutes post HFS. All post-hoc pairwise comparisons reported are Sidak-Holm corrected.

**Table 2.** ANOVA results from percentage of change in MPS measures. Significance from post-hoc pairwise comparisons are reported following Sidak Holm multiple comparisons correction. rrmm = repeated measures; HFS = high frequency stimulation.

| Test | Main effects |  |  |  | Pairwise comparisons |  |  |
| --- | --- | --- | --- | --- | --- | --- | --- |
|  | Condition & Time |  | Interaction effect |  | Mean difference (*p*) |  |  |
|  | F(df) | *P* | F(df) | *p* | PreHFS-15post | PreHFS-35post | PreHFS-50post |
|  |  |  |  |  | HFS | HFS | HFS |
| Test arm |  |  |  |  |  |  |  |
|  | C:0.092<sub>(2,56)</sub> | 0.859 |  |  |  |  |  |
| Two-way rrm ANOVA | T:6.098<sub>(2,56)</sub> | 0.012 | 0.147<sub>(6,168)</sub> | 0.812 | - | - | - |
| One-way ANOVA – no intervention | 8.495<sub>(3,84)</sub> | <0.001 | - | - | -159.19<br>(0.019) | -169.52<br>(<0.001) | -136.84<br>(0.001) |
| One-way ANOVA – VR | 3.477<sub>(3,84)</sub> | 0.060 | - | - | -205.09<br>(0.320) | -196.28<br>(0.209) | -163.09<br>(0.450) |
| One-way ANOVA – sham VR | 2.339<sub>(3,84)</sub> | 0.133 | - | - | -190.14<br>(0.714) | -205.28<br>(0.512) | -201.76<br>(0.254) |
| Control arm |  |  |  |  |  |  |  |
|  | C:0.906<sub>(2,56)</sub> | 0.410 |  |  |  |  |  |
| Two-way rrm ANOVA | T:1.300<sub>(2,56)</sub> | 0.279 | 0.783<sub>(6,168)</sub> | 0.434 | - | - | - |
| One-way ANOVA – no intervention | 1.593<sub>(3,84)</sub> | 0.218 | - | - | -85.15<br>(0.668) | -63.40<br>(0.899) | -92.96<br>(0.699) |
| One-way ANOVA – VR | 3.000<sub>(3,84)</sub> | 0.047 | - | - | 17.85<br>(0.693) | 25.79<br>(0.045) | -9.09<br>(0.969) |
| One-way ANOVA – sham VR | 0.706<sub>(3,84)</sub> | 0.413 | - | - | -115.36<br>(0.940) | -32.80<br>(0.981) | -31.14<br>(0.578) |

*ΔMPS:* Within the test arm, two-way repeated measures ANOVA revealed a main effect of Condition (F_(2,56)_=5.234, *p*=0.013), a main effect of Timepoint (F_(3,84)_=18.322, *p*<0.001) and a significant Condition x Timepoint interaction effect (F_(6,168)_=3.571, *p*=0.021). Within each condition, one-way repeated measures ANOVA’s showed a significant effect of Time across all conditions (no intervention: F_(3,84)_=8.86, *p*<0.001; sham-VR: F_(3,84)_=8.86, *p*<0.001; VR: F_(3,84)_=5.206, *p*=0.013). Post-hoc pairwise comparisons revealed that in the no intervention condition, ΔMPS were significantly different from baseline at all timepoints post HFS (Figure 3). On the sham-VR condition, ΔMPS was significantly greater than baseline at 35- and 50-minutes post HFS. On the VR condition, pairwise comparisons revealed no significant differences across all timepoints post HFS (Table 3). Within each individual pinprick intensity, analyses yielded almost identical results; however, within the 128mN prinprick stimuli there was no longer a significant MPS increase from baseline at 35-minutes and 50-minutes post HFS in the sham-VR condition, and there was a significant MPS increase from baseline at 15-minutes post HFS during the VR intervention.

**Figure 3.**
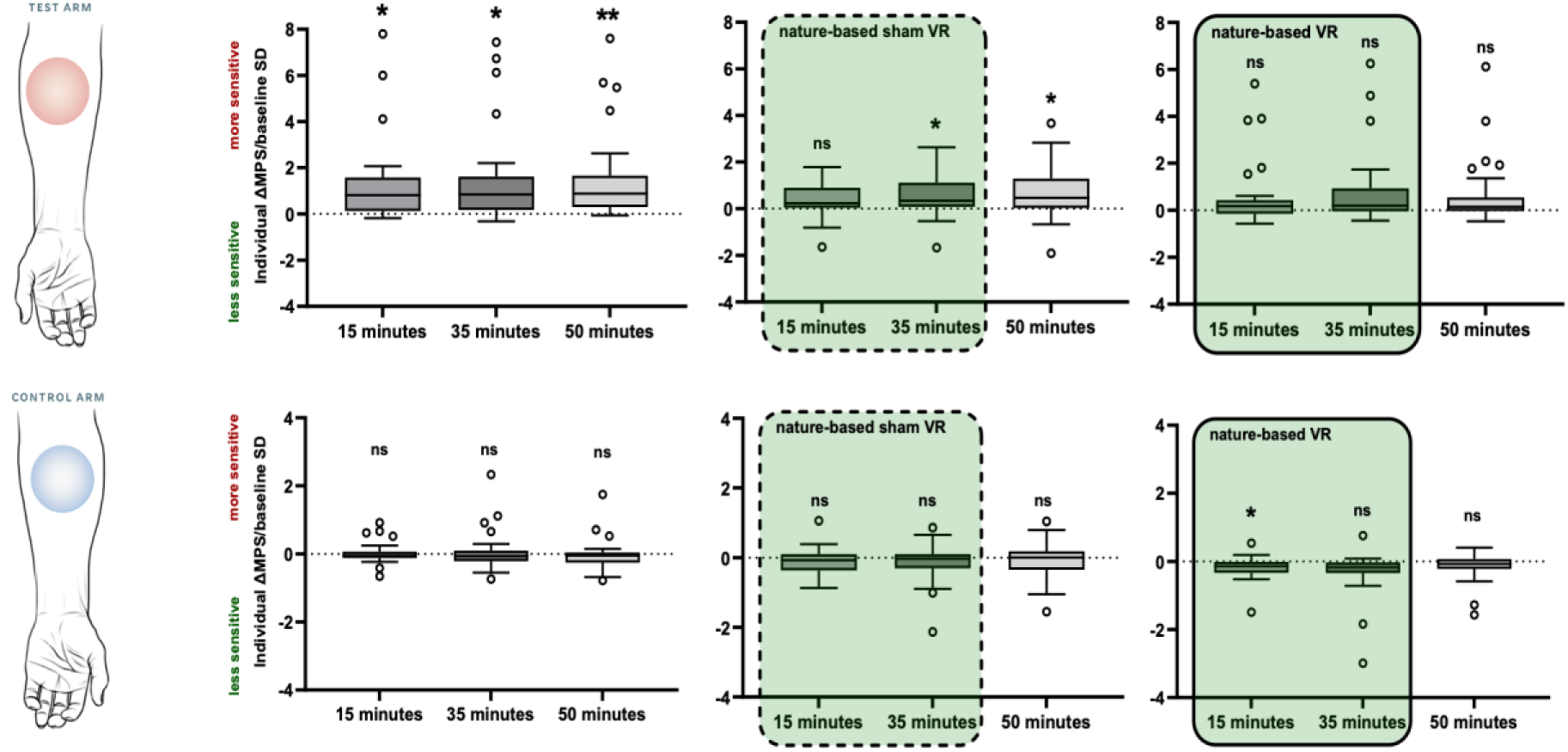
MPS results for the test arm (top row) and the control arm (bottom row). Boxplots represent changes in MPS from baseline during the no intervention session (left graphs), during the sham VR session (middle graphs) and during the VR session (right graphs). Sham VR and VR interventions started immediately following HFS and terminated at ∼45mins post-HFS. All results are depicted as standard measures (i.e., changes in MPS form baseline divided by standard deviation of baseline MPS results within each session). *Significant at alpha 0.05; **Significant at alpha 0.01; ns= not significant.

**Table 3.**
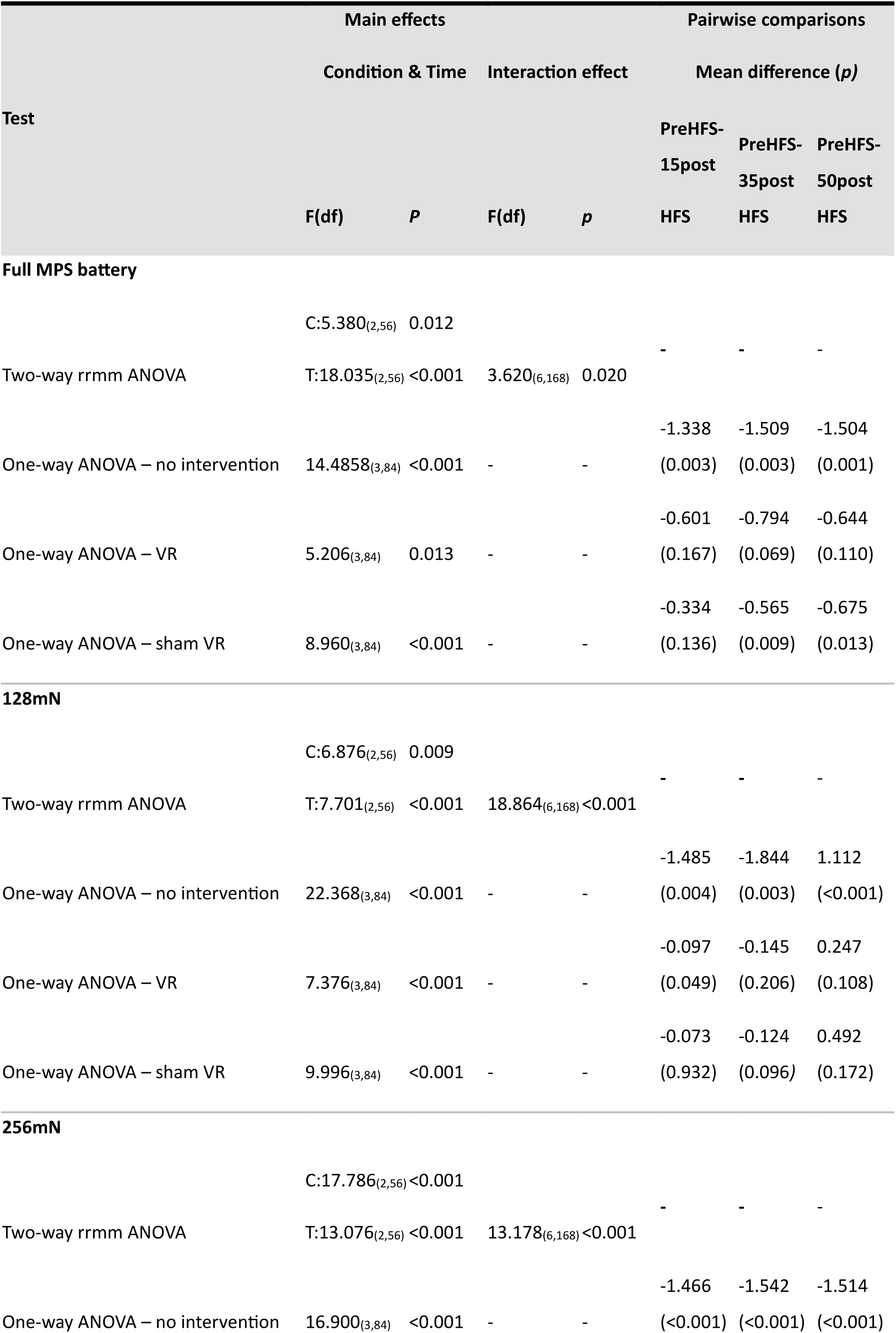
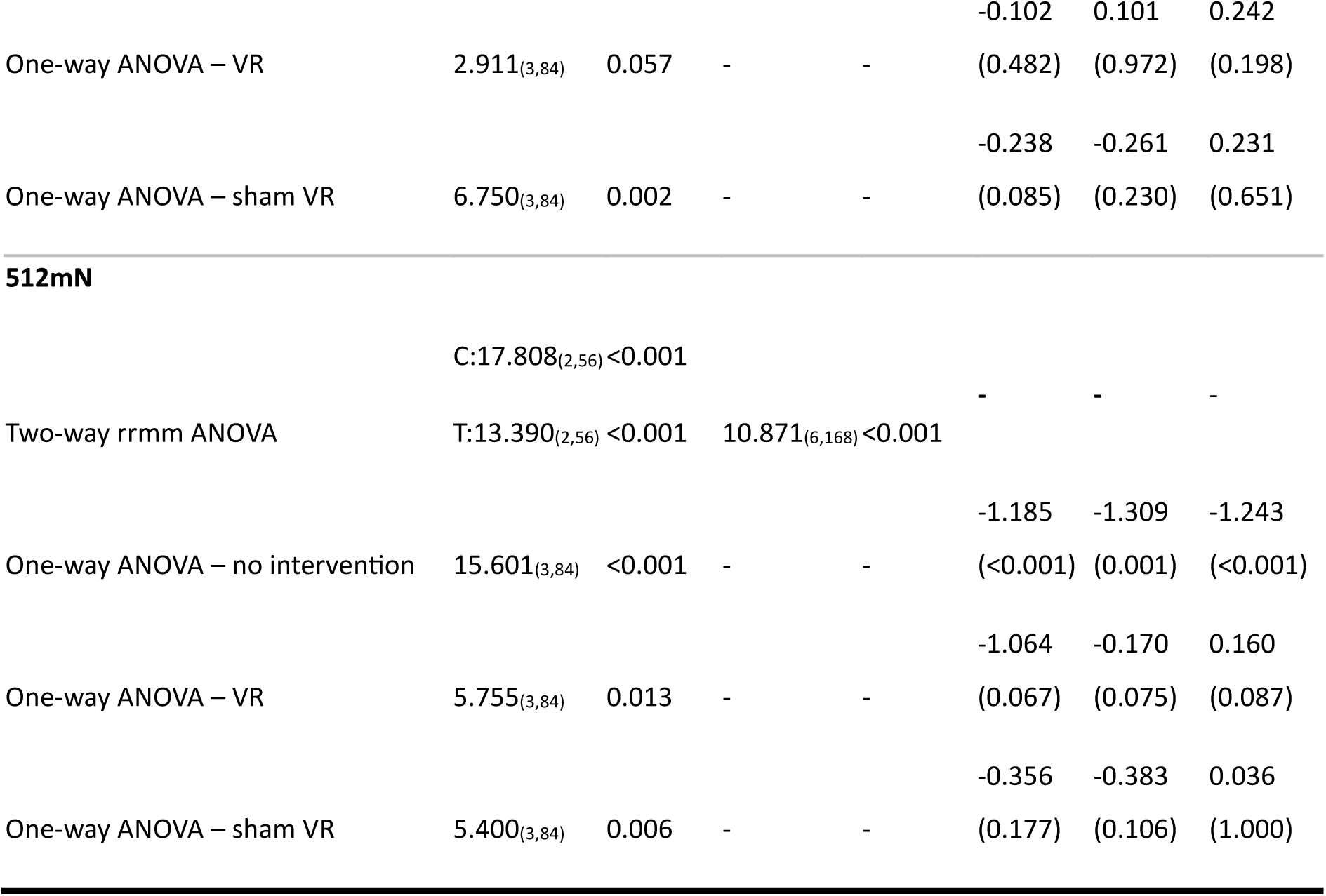
ANOVA results from standard change in MPS measures from baseline (test arm). Significance from post-hoc pairwise comparisons are reported following Sidak Holm multiple comparisons correction. rrmm = repeated measures; HFS = high frequency stimulation.

| Test | Main effects |  |  |  | Pairwise comparisons |  |  |
| --- | --- | --- | --- | --- | --- | --- | --- |
|  | Condition & Time |  | Interaction effect |  | Mean difference (*p*) |  |  |
|  | F(df) | *P* | F(df) | *p* | PreHFS-15post | PreHFS-35post | PreHFS-50post |
|  |  |  |  |  | HFS | HFS | HFS |
| Full MPS battery |  |  |  |  |  |  |  |
|  | C:5.380<sub>(2,56)</sub> | 0.012 |  |  | - | - | - |
| Two-way rrrmm ANOVA | T:18.035<sub>(2,56)</sub> | <0.001 | 3.620<sub>(6,168)</sub> | 0.020 |  |  |  |
| One-way ANOVA – no intervention | 14.4858<sub>(3,84)</sub> | <0.001 | - | - | -1.338<br>(0.003) | -1.509<br>(0.003) | -1.504<br>(0.001) |
| One-way ANOVA – VR | 5.206<sub>(3,84)</sub> | 0.013 | - | - | -0.601<br>(0.167) | -0.794<br>(0.069) | -0.644<br>(0.110) |
| One-way ANOVA – sham VR | 8.960<sub>(3,84)</sub> | <0.001 | - | - | -0.334<br>(0.136) | -0.565<br>(0.009) | -0.675<br>(0.013) |
| 128mN |  |  |  |  |  |  |  |
|  | C:6.876<sub>(2,56)</sub> | 0.009 |  |  | - | - | - |
| Two-way rrrmm ANOVA | T:7.701<sub>(2,56)</sub> | <0.001 | 18.864<sub>(6,168)</sub> | <0.001 |  |  |  |
| One-way ANOVA – no intervention | 22.368<sub>(3,84)</sub> | <0.001 | - | - | -1.485<br>(0.004) | -1.844<br>(0.003) | 1.112<br>(<0.001) |
| One-way ANOVA – VR | 7.376<sub>(3,84)</sub> | <0.001 | - | - | -0.097<br>(0.049) | -0.145<br>(0.206) | 0.247<br>(0.108) |
| One-way ANOVA – sham VR | 9.996<sub>(3,84)</sub> | <0.001 | - | - | -0.073<br>(0.932) | -0.124<br>(0.096) | 0.492<br>(0.172) |
| 256mN |  |  |  |  |  |  |  |
|  | C:17.786<sub>(2,56)</sub> | <0.001 |  |  | - | - | - |
| Two-way rrrmm ANOVA | T:13.076<sub>(2,56)</sub> | <0.001 | 13.178<sub>(6,168)</sub> | <0.001 |  |  |  |
| One-way ANOVA – no intervention | 16.900<sub>(3,84)</sub> | <0.001 | - | - | -1.466<br>(<0.001) | -1.542<br>(<0.001) | -1.514<br>(<0.001) |
|  |  |  |  |  | -0.102 | 0.101 | 0.242 |
| One-way ANOVA – VR | 2.911<sub>(3,84)</sub> | 0.057 | - | - | (0.482) | (0.972) | (0.198) |
|  |  |  |  |  | -0.238 | -0.261 | 0.231 |
| One-way ANOVA – sham VR | 6.750<sub>(3,84)</sub> | 0.002 | - | - | (0.085) | (0.230) | (0.651) |
| **512mN** |  |  |  |  |  |  |  |
|  | C:17.808<sub>(2,56)</sub> | <0.001 |  |  | - | - | - |
| Two-way rmm ANOVA | T:13.390<sub>(2,56)</sub> | <0.001 | 10.871<sub>(6,168)</sub> | <0.001 |  |  |  |
|  |  |  |  |  | -1.185 | -1.309 | -1.243 |
| One-way ANOVA – no intervention | 15.601<sub>(3,84)</sub> | <0.001 | - | - | (<0.001) | (0.001) | (<0.001) |
|  |  |  |  |  | -1.064 | -0.170 | 0.160 |
| One-way ANOVA – VR | 5.755<sub>(3,84)</sub> | 0.013 | - | - | (0.067) | (0.075) | (0.087) |
|  |  |  |  |  | -0.356 | -0.383 | 0.036 |
| One-way ANOVA – sham VR | 5.400<sub>(3,84)</sub> | 0.006 | - | - | (0.177) | (0.106) | (1.000) |

Within the control arm (Table 4), we observed no significant main effects of Condition or Timepoint (F_(2,56)_=2.480, *p*=0.095 and F_(3,84)_=1.644, *p*=0.209, respectively) There was a significant Condition x Timepoint interaction effect (F_(6,168)_=2.691, *p*=0.046). In the VR condition, there was a significant effect of Time (F_(3,84)_=4.843, *p*=0.020), driven by a significant reduction in MPS at 15-minutes post HFS compared to baseline (Figure 3). There were no significant effects of Time for the sham-VR or the no intervention conditions (F_(3,84)_=1.121, *p*=0.336 and F_(3,84)_=0.410, *p*=0.652, respectively). Within each pinprick intensity, we observed significant main effects of Condition and Timepoint. One-way ANOVA’s revealed main effects of Timepoint for VR and sham-VR interventions (Table 4). Post-hoc pairwise comparisons indicated that there was a significant MPS decrease from baseline at 15-minutes post HFS during the VR intervention for the 128mN pinprick but not for the others. Individual pinprick results also displayed a significant MPS decrease post sham-VR intervention compared to baseline.

**Table 4.** ANOVA results from standard change in MPS measures from baseline (control arm). Significance from post-hoc pairwise comparisons are reported following Sidak Holm multiple comparisons correction. rrmm = repeated measures; HFS = high frequency stimulation.

| Test | Main effects |  |  |  | Pairwise comparisons |  |  |
| --- | --- | --- | --- | --- | --- | --- | --- |
|  | Condition & Time |  | Interaction effect |  | Mean difference (*p*) |  |  |
|  | F(df) | *P* | F(df) | *p* | PreHFS- | PreHFS- | PreHFS- |
|  |  |  |  |  | 15postHFS | 35postHFS | 50postHFS |
| Full MPS battery |  |  |  |  |  |  |  |
|  | C:2.328<sub>(2,56)</sub> | 0.109 |  |  |  |  |  |
| Two-way rrm ANOVA | T:1.696<sub>(2,56)</sub> | 0.200 | 2.768<sub>(6,168)</sub> | 0.038 | - | - | - |
|  |  |  |  |  | -0.027 | -0.059 | 0.027 |
| One-way ANOVA – no intervention | 0.410<sub>(3,84)</sub> | 0.652 | - | - | (0.998) | (0.995) | (1.000) |
|  |  |  |  |  | 0.185 | 0.309 | 0.143 |
| One-way ANOVA – VR | 4.843<sub>(3,84)</sub> | 0.020 | - | - | (0.036) | (0.097) | (0.368) |
|  |  |  |  |  | 0.122 | 0.134 | 0.067 |
| One-way ANOVA – sham VR | 1.121<sub>(3,84)</sub> | 0.038 | - | - | (0.550) | (0.729) | (0.982) |
| 128mN |  |  |  |  |  |  |  |
|  | C:6.876<sub>(2,56)</sub> | 0.009 |  |  |  |  |  |
| Two-way rrm ANOVA | T:7.701<sub>(2,56)</sub> | <0.003 | 18.864<sub>(6,168)</sub> | <0.001 | - | - | - |
|  |  |  |  |  | -0.052 | 0.002 | 0.121 |
| One-way ANOVA – no intervention | 0.489<sub>(3,84)</sub> | 0.572 | - | - | (0.998) | (1.000) | (0.990) |
|  |  |  |  |  | 0.815 | -0.063 | -0.010 |
| One-way ANOVA – VR | 15.136<sub>(3,84)</sub> | <0.001 | - | - | (<0.001) | (0.897) | (0.183) |
|  |  |  |  |  | 0.012 | 0.020 | 1.074 |
| One-way ANOVA – sham VR | 25.042 | <0.001 | - | - | (1.000) | (1.000) | (<0.001) |
| 256mN |  |  |  |  |  |  |  |
|  | C:14.142<sub>(2,56)</sub> | <0.001 |  |  |  |  |  |
| Two-way rrm ANOVA | T:11.197<sub>(2,56)</sub> | <0.001 | 20.084<sub>(6,168)</sub> | <0.001 | - | - | - |
|  |  |  |  |  | -0.022 | 0.043 | 0.000 |
| One-way ANOVA – no intervention | 0.137<sub>(3,84)</sub> | 0.872 | - | - | (1.000) | (0.996) | (1.000) |
|  |  |  |  |  | -0.022 | 0.043 | 0.000 |
| One-way ANOVA – VR | 3.121<sub>(3,84)</sub> | 0.049 | - | - | (1.000) | (0.996) | (1.000) |
|  |  |  |  |  | 0.078 | 0.039 | 1.124 |
| One-way ANOVA – sham VR | 31.845 | <0.001 |  |  | (0.717) | (0.998) | (<0.001) |
| **512mN** |  |  |  |  |  |  |  |
|  | C:4.807<sub>(2,56)</sub> | 0.021 |  |  |  |  |  |
| Two-way rrm ANOVA | T 6.626<sub>(2,56)</sub> | 0.004 | 23.520<sub>(6,168)</sub> | <0.001 | - | - | - |
|  |  |  |  |  | 0.045 | 0.102 | 0.072 |
| One-way ANOVA – no intervention | 0.353<sub>(3,84)</sub> | 0.651 | - | - | (0.992) | (0.451) | (0.994) |
|  |  |  |  |  | 0.254 | -0.025 | -0.151 |
| One-way ANOVA – VR | 2.891<sub>(3,84)</sub> | 0.064 | - | - | (0.401) | (0.999) | (0.173) |
|  |  |  |  |  | -0.080 | - | 1.111 |
| One-way ANOVA – sham VR | 34.735<sub>(3,84)</sub> | 0.001 | - | - | (0.977) | 0.132(0.335) | (<0.001) |

#### Secondary hyperalgesia mapping

Within the test arm, participants rated stimuli across all eight axes as more painful than at baseline, consistently across all post HFS timepoints (Figure 4) during the no intervention session. Peaks of hyperalgesia mapping were observed along the proximodistal direction, extending laterally along axes 2 and 6. During the sham-VR session, the magnitude of hyperalgesia was reduced across the mapping area across all timepoints post HFS, with a peak of moderate hyperalgesia around 1cm away from the electrode in the laterodistal direction (axis 6). In the VR session, no peaks reaching moderate hyperalgesia were identified. Within the control arm, we observed no traces of hyperalgesia across conditions.

**Figure 4.**
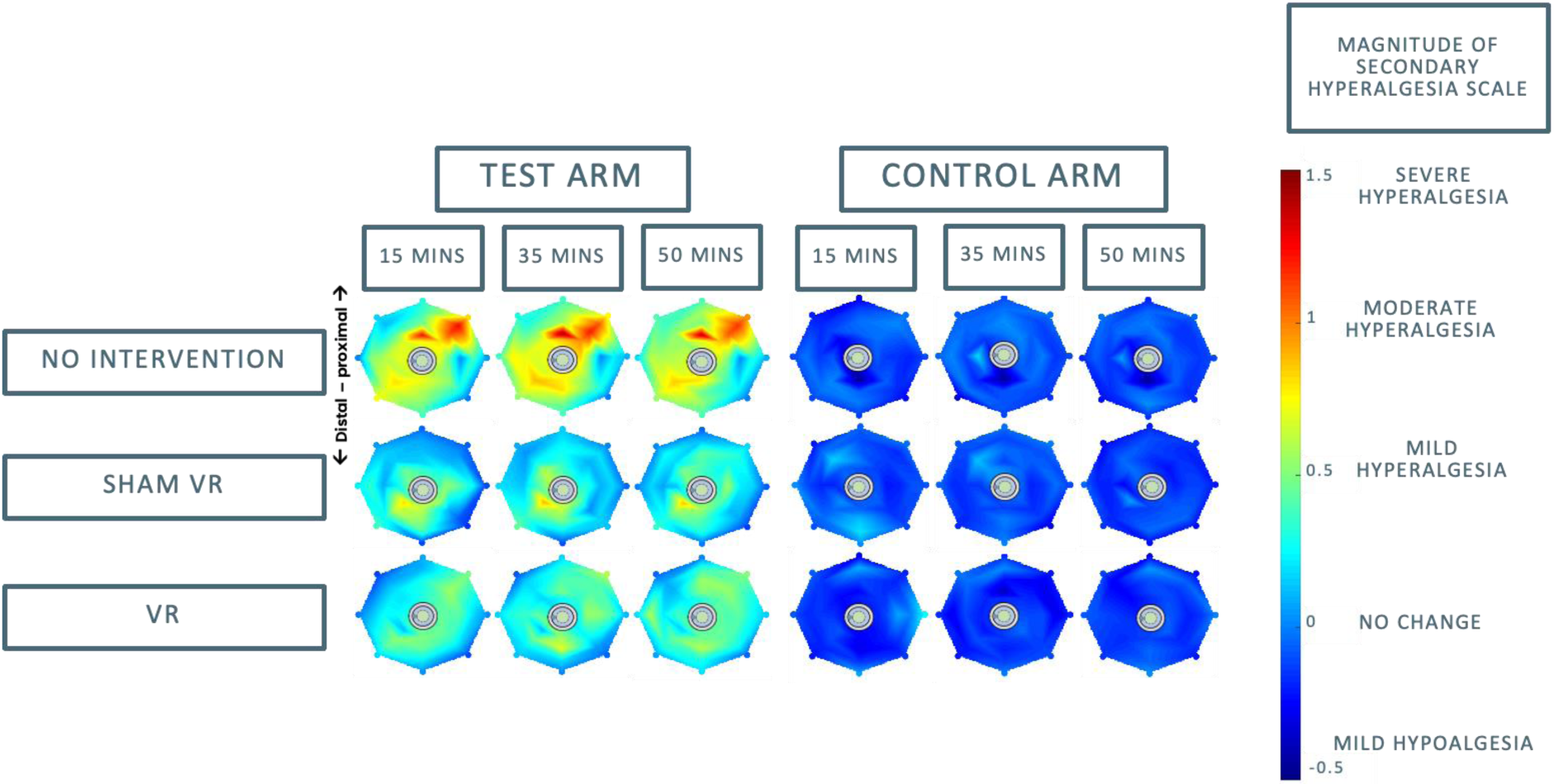
Secondary hyperalgesia mapping results. Each colourmap was calculated by interpolating results across adjacent mapping points across all axes (last mapping point for each axis is depicted as a dot by the edge of the map). Grey circles in the middle of each map denote the location of the HFS electrode (this was drawn for both arms, but no HFS was performed on the control arm). Measures of mild, moderate and severe hyperalgesia were arbitrarily assigned standard changes in NRS from baseline (i.e., absolute changes from baseline on each landmark divided by standard deviation of baseline NRS across the group) equal to 0.5, 1 and 1.5, respectively.

**Figure 5.**
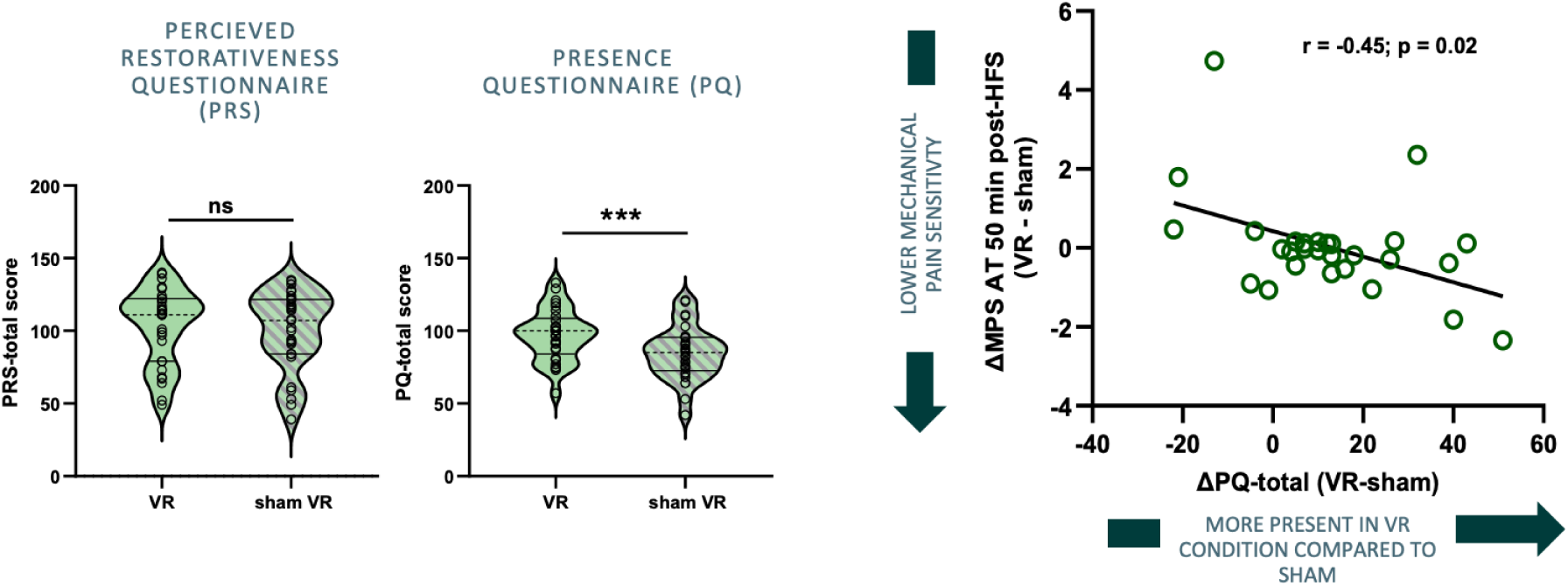
Results from psychometric assessment of VR intervention experience. Results from paired-samples t-test within PRS and PQ scales are depicted on the left. There was no significant difference across VR and sham-VR sessions for the PRS, however PQ scores were significantly higher in the VR condition. Scatter plot on the right shows Pearson’s correlation results between difference in delta MPS 50 mins post-HFS (VR – sham) and difference in PQ results (VR-sham). Negative correlation indicates that participants who reported higher levels of presence during the VR intervention compared to sham also experience lower hyperalgesia following VR intervention.

### Relationship between individual experience during interventions and differential analgesic effects post intervention

Pearson’s correlation analysis yielded a significant negative relationship between delta 50-minutes post HFS MPS scores (VR – sham) and delta PQ scores (VR – sham), (r=-0.45, *p*=0.02). This is, participants reporting higher levels of presence during the VR intervention compared to the sham-VR intervention also reported lower levels of mechanical pain sensitivity following VR intervention than following sham-VR. There was no significant correlation between delta PRS scores and delta 50-minutes post HFS MPS scores.

### Spectral DCM results

#### Accuracy of the DCM estimation

The mean variance (SD) explained by DCM model estimations when fitted to the observed spectral data was 90.5% (2.5%), indicating successful estimation of DCMs to the observed timeseries.

#### Commonalities of effective connectivity during tonic noxious stimulation across all participants

Results from greedy search over the first 256 reduced models indicated that there was not an overall winning model (all ∼1% probability), this is, the exclusion of parameters during the nesting process did not lead to a substantial improvement in model fit. However, BMA results for the group mean effect indicated that several parameters had a 95% probability of being present vs absent (Figure 6b). Specifically, we observed lateralised increased self-inhibition across right dACC, right AI and right thalamus, and decreased self-inhibition in the left thalamus and PAG. We observed overall excitatory effective connectivity in the top-down direction as well as inhibitory effective connectivity in the bottom-up direction.

**Figure 6.**
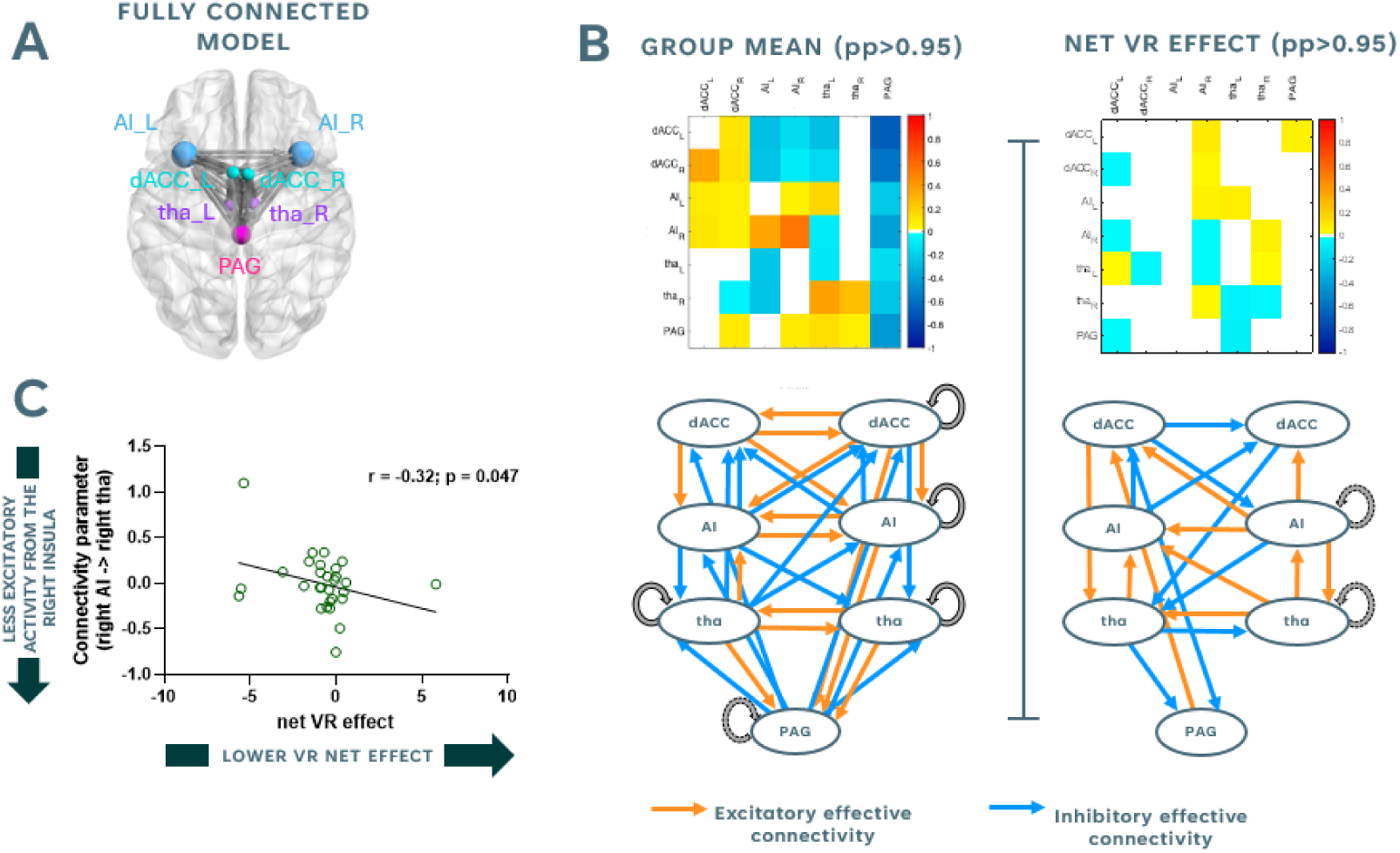
Summary of spDCM results. (A) Locations of seven nodes comprising the descending pain control pathway used for spDCM analysis. Due to the exploratory nature of the analysis, a fully connected model was initially described. (B) Results from greedy search through nested models using Bayesian model reduction (BMR). Left panel shows commonalities across all subjects during tonic pain. Right panel shows continues linear relationship between effective connectivity and net VR effect (i.e., delta MPS at 50 mins post-HFS during VR conditions - delta MPS at 50 mins post-HFS during baseline conditions). Off-diagonal positive numbers (represented by orange arrows on the corresponding bottom diagram) represent excitatory connections (*from* locations indicated by rows and *to* locations indicated by columns), and off-diagonal negative numbers (represented by blue arrows on the corresponding bottom diagram) represent inhibitory connections. (C) Pearson’s correlation between strength of excitatory connectivity form right AI to right thalamus (selected from CVA results as parameter with highest load on the canonical variate) and net VR effect. AI_L = left anterior insula; AI_R = right anterior insula; dACC_L = left dorsal anterior cingulate cortex; dACC_R = right dorsal anterior cingulate cortex; tha_L = left thalamus; tha_R = right thalamus; PAG = periaqueductal grey; pp = posterior probability.

#### Relationship between VR intervention effects and autonomic descending pain regulation network architecture

Parameters with posterior probabilities >95% for the continuous linear relationship between the defined network and net VR effect exhibited, in contrast, decreased self-inhibition in right AI and right thalamus. We also observed a switch in the direction of excitatory/inhibitory connections, with inhibitory connectivity primarily directed top-down and excitatory activity primarily observed for top-up connections.

#### Correlation with nature-VR pain measures

The connectivity parameter for the connection from right AI to right thalamus exhibited the highest loading on the canonical variate (0.66). In order to better interpret its relationship with the net VR effects, we computed the one-tailed Pearson’s correlation between the ‘right AI –> right thalamus’ connectivity strength parameter across participants and the net VR effects (Figure 6c), revealing a significant negative correlation (r=-0.32, *p*=0.047). This is, participants exhibiting higher excitatory effective connectivity from the right AI to the right thalamus during tonic pain tended to have a greater benefit from the VR intervention compared to baseline.

## Discussion

In this study, we have demonstrated that a 45-minute immersive nature experience delivered through VR can attenuate both the development and heterotopic spread of mechanical secondary hyperalgesia in healthy participants. VR nature experiences induced a higher sense of presence compared to non-immersive nature experiences, which also had weaker, non-lasting analgesic effects on mechanical secondary hyperalgesia. Additionally, top-down insulo-thalamic effective connectivity during noxious stimulation related to greater benefits from VR nature immersion in the same individuals, further highlighting the potentially direct effects of nature on endogenous pain modulatory systems.

HFS can successfully induced sustained mechanical secondary hyperalgesia as early as 15 minutes post-stimulation in the no intervention condition. Given that the development of secondary hyperalgesia is thought to be a manifestation of central sensitisation and is a key feature of many chronic pain states [26], it serves as a biomarker to help understand the mechanisms of nature-based VR. We demonstrated that prolonged virtual immersion in nature effectively produced analgesic effects during the development of mechanical secondary hyperalgesia, evidenced by the lack of increase in mechanical pain sensitivity observed during -and following-exposure to nature.

Several factors should be considered to interpret the effects of nature VR on the development of mechanical secondary hyperalgesia; firstly, these are unlikely to be attributed to natural variability in MPS measures across sessions, as MPS has been shown to reliably capture HFS-induced chances in pain sensitivity on different test days [27]. Secondly, while disentangling the analgesic effects of nature from pure distraction caused by VR poses inherent challenges [28], current distraction-based VR interventions for pain typically required specific cognitive strategies or games [29, 30] to achieve benefits. In chronic pain populations, these are often limited to improving coping mechanisms rather than reducing pain directly [31]. In addition, previous attempts to modulate the development of experimentally induced mechanical secondary hyperalgesia using non-VR based working memory manipulations delivered during the conditioning period have produced mixed results [32, 33]. Moreover, distraction-based strategies can exacerbate pain following distraction termination [34]. This does not align with our approach, as we used a 45-minute immersive *and passive* nature-based VR intervention given immediately after the conditioning period, to co-inside with the development of secondary hyperalgesia. Participants were also instructed to focus on their arms during stimulation across all sessions, and analgesic effects persisted for at least five minutes post-VR intervention. It is plausible, however, that distraction influenced the pain sensitivity measures in the sham condition. The cognitive and attention demands of watching a video on a screen may have prevented a significant rise in hypersensitivity measures initially, but as the novelty of the scene diminished, secondary hyperalgesia eventually developed.

A key finding supporting the existence of differential mechanisms modulating secondary hyperalgesia in the sham and VR conditions is the significant increase in perceived sense of presence during the VR intervention. It is important to emphasise that presence and distraction as distinct concepts. In this context, presence refers to the feeling of being fully immersed in a virtual environment as if it were the real world, without necessarily directing attention to or away from specific aspects [35]. In contrast, distraction involves the redirection of attention away from the environment or oneself, without requiring immersion of any sort. Critically, participants who reported a greater sense of presence during VR compared to sham also experience greater analgesic effects following the intervention. Taken together, our results suggest that a 2D nature scene on a screen might not be sufficient to elicit therapeutic effects. However, achieving high levels of presence in a natural environment through VR may more closely replicate the benefits of real nature immersion. This has important implications for the use of virtual nature in social prescribing for chronic pain, as our findings indicate that a prolonged nature-based VR intervention could potentially capture the positive elements of nature while mitigating the challenges, such as inaccessibility, the risk of injury or time constraints.

A secondary aim of this study was to shed light into the top-down pain-related mechanisms by which nature scenes may influence the development of sensitised pain states. To help address this question, we introduced a novel, more nuanced approach to mapping the area of secondary hyperalgesia following HFS. While previous methods relied on assessing the presence or absence of pain at various points along proximodistal axes of the HFS site [36, 37], we employed NRS ratings before and after HFS along radial spokes in order to determine the magnitude of hyperalgesia that spreads across the heterotopic area. This approach demonstrated that as well producing temporal effects, nature-based VR can also spatially restrict the spread of HFS-induced secondary hyperalgesia. Given that the extent of heterotopic spread of pin prick sensitivity is by definition a form of spinal plasticity, it is possible to use these psychophysical data as a proxy for investigating descending influences on spinal nociception. Given that nature-based analgesia is thought to engage a combination of both autonomic and cognitive influences, it is therefore feasible that these cortical mechanisms could be activating top-down spinally-projecting pathways.

In fact, we found that reduced excitatory influence from the AI to the thalamus during a painful state was associated with weaker analgesic effects of nature immersion. This finding is particularly compelling, given the critical role of both regions in modulating autonomic responses. Recent metanalytical evidence from neuroimaging studies indicate that the AI plays a critical role in the central autonomic system [38]; the AI integrates current interoceptive states, stimulus intensity and cognitive processing, and can influence autonomic regulation dynamically via its projections to the hypothalamus, the amygdala and basal ganglia [39]. The thalamus is a key hub for gathering information on sympathetic and parasympathetic tone from the periphery directly, as well as via brainstem nuclei [38], which serves to update interoceptive states [40]. This monitoring includes pain-related inputs via spino-thalamic-cortical pathways that project to the AI, unsurprisingly making AI dysregulation a potentially key functional biomarker of chronic pain [41]. We may therefore speculate that individuals experiencing greater analgesic effects from immersion in nature in our study were those who spontaneously recruited top-down insulo-thalamic pain regulatory pathways during pain more optimally. Taken together, our results represent further, novel evidence supporting the role if nature in pain modulation via brain regions involved in autonomic regulation. While these are undoubtedly encouraging results, more research into the exact cortical, sub-cortical and brainstem mechanisms alongside physiological measures are required in order to fully interpret these results.

In contrast to our a priori hypothesis, we observed no significant differences in perceived restorativeness between VR and sham interventions. Although initially surprising, it is key to consider that participants may have provided responses based on how they *would feel* in the environment, rather than reflecting their experience in the moment. Therefore, the PRS may not have been the most sensitive tool for capturing the immediate sense of restoration during VR. Future studies with similar designs may benefit from employing more tailored methods for the assessment of perceived restorativeness in a VR environment.

Our study also presented some limitations. Our spDCM did not reach the same sample size as our main analyses, which might have hampered the statistical power of our results. This could also have influenced the fact that although we observed specific patterns of connections with very high probability of being present and were highly informative, we did not find a single model that was superior in explaining variability in nature effects. It is possible that a different outcome would have been obtained if including lower midbrain and brainstem nuclei in our model had been a possibility, however our reduced field of view (necessary to keep optimal temporal and spatial resolution) did not allow for this to occur. Our results, however, can serve to guide future efforts utilising a similar approach, where more complex models more specific to autonomic regulation may be tested.

In conclusion, our findings provide novel critical insights into the analgesic mechanisms associated with prolonged exposure to immersive nature-based environments, further supporting the integration of nature into social prescribing for pain management. By demonstrating that a 45-minute immersive nature VR experience can attenuate the development and spread of mechanical secondary hyperalgesia and revealing a relationship between effective AI – insula effective connectivity with enhanced analgesic effects, we provide critical insights into the top-down endogenous analgesic mechanisms involved. These results suggest that immersive nature experiences can modulate pain through brain regions involved in autonomic pain modulation which could be harnessing endogenous analgesic activity in spinally-projecting descending pain modulation systems. Future research should focus on elucidating detailed physiological, brainstem and spinal mechanisms of nature-based analgesia and should translate these findings into novel pain management strategies for chronic pain.

## Acknowledgements

This study was funded through an Academy of Medical Sciences Springboard grant (SBF007\100108). We would like to thank Alexander Smith for his input and help to select the nature footage. We would like to express our immense gratitude to all the participants that volunteered for this study.

