## Supplementary Information for "Immersion in nature attenuates the development of mechanical secondary hyperalgesia: a role for insulo-thalamic effective connectivity"

### a. MRI session

Procedures

MRI scanning was conducted at the Mireille Gillings Neuroimaging Centre in Exeter. Prior to the session, participants' adherence to lifestyle guidelines and MRI eligibility were reviewed. All participants completed the state portion of the State Trait Anxiety Inventory (STAI) [1] to ensure no clinically significant levels of anxiety were reached on the day of the scan. The session began with a localizer and structural scan, followed by three resting-state functional magnetic resonance imaging (rs-fMRI) scans, each lasting about six minutes. The first rs-fMRI scan served as a baseline to evaluate brain function at rest without external stimulation. The second scan represented the tonic pain condition; before it started, a cold gel tub was placed on the participants' abdomen over a plastic sheet to prevent direct cooling of the skin. Participants were instructed to fully immerse their right hand into the cold gel at the start of the scan and keep it still throughout. After the scan ended, they removed their hand, and the experimenter took away the gel tub. The third scan replicated the baseline scan to measure brain function during recovery from tonic pain. At the start of the baseline scan and at the end of each subsequent scan, participants rated their current pain on a numerical rating scale (NRS) from 0 (no pain) to 100 (worst imaginable pain). After the tonic pain scan, they also rated their average pain during the cold stimulation on an NRS. Throughout all rs-fMRI scans, participants were instructed to lie still with their eyes open, focusing on a white fixation cross on a black background displayed on a screen in front of them.

Materials

The method for inducing tonic cold pain followed the procedure of Lapotka et al. (2017) [2]. The gel consisted of a mixture of 300g cornstarch, 2 litres of water, and 100g salt, prepared in a standard 2-liter rectangular plastic container. Pilot tests showed that the sub-1°C temperatures used by Lapotka et al. were intolerable for most pilot participants during the full scan duration, therefore tubs were refrigerated at 5°C for at least 24 hours before each session. This allowed for tolerable six-minute stimulation that still produced varying levels of tonic pain. To maintain a consistent initial te[3]mperature, the tubs were removed from the fridge no more than two minutes before being used in the scanner room or presented after the scan. Fresh tubs were used for each hand during the session.

Data preprocessing

For fMRI data preprocessing, motion correction was applied using MCFLIRT [4] from FSL [5] to address head movements. Geometric distortions due to magnetic field inhomogeneities were corrected using the FUGUE command in FSL FEAT. Multi-echo fMRI data were denoised with TEDANA [6], employing principal component analysis (PCA) and independent component analysis (ICA) for denoising. Signals from white matter (WM) and cerebrospinal fluid (CSF) were regressed out using masks derived from T1-weighted scans segmented with DARTEL in SPM12 [3, 7] and coregistered to functional scans with FLIRT. The data were then high-pass filtered at 0.005 Hz to remove low-frequency drifts. Normalisation to MNI space involved linear and non-linear warping of grey matter (GM) masks, followed by application of these warping parameters to the functional images. Finally, spatial smoothing was conducted with a 5mm FWHM Gaussian kernel to enhance the signal-to-noise ratio.

### b. ROI definition

The regions of interest (ROIs) for the dorsal anterior cingulate cortex (dACC), anterior insula (AI), and thalamus were delineated using spherical masks with centers at MNI coordinates derived from probabilistic maps in the Brainnetome Atlas. Specific map labels, MNI coordinates, and radius sizes are detailed in the table below. For the periaqueductal gray (PAG), anatomical masks were obtained from the Brainstem Navigator toolkit.

| ROI | BNA atlas labels | MNI coordinates | | Radius size |
|---|---|---|---|---|
| | | Left | Right | |
| dorsal ACC | A24cd, caudodorsal area 24 | -5, 7, 37 | 4, 6, 38 | 4mm |
| AI | dIa, dorsal agranular insula | -34, 18, 1 | 36, 18, 1 | 8mm |
| thalamus | mPFtha, medial pre-frontal thalamus | -7, -12, 5 | 7, -11, 6 | 4mm |

**Supplementary table 1.** ROI definition details. ACC = anterior cingulate cortex; AI = anterior insula; MNI = Montreal Neurological Institute

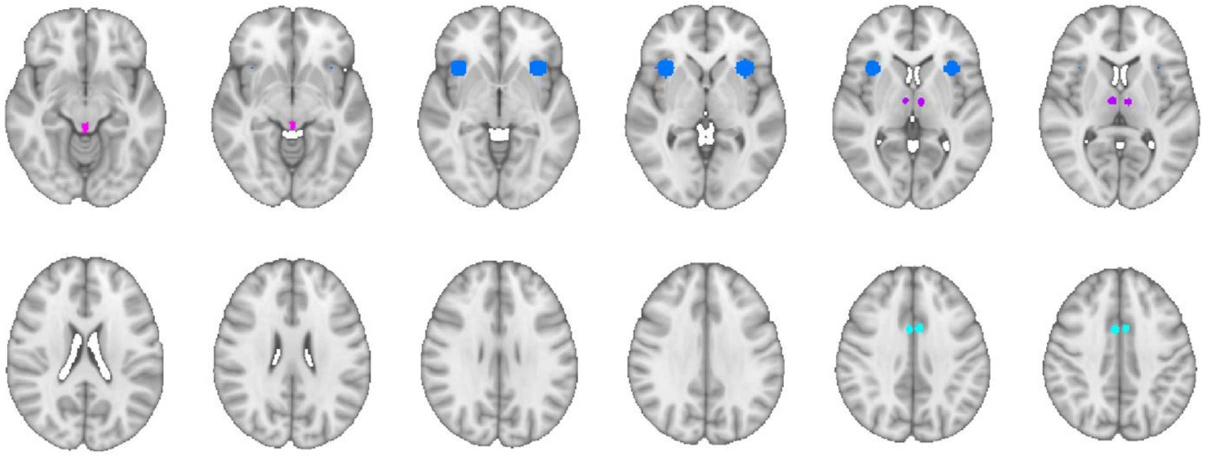

**Supplementary Figure 1.** Depiction of dACC (light blue), AI (dark blue), thalamus (purple) and PAG (pink) ROIs.
